# Aequatus: An open-source homology browser

**DOI:** 10.1101/055632

**Authors:** Anil S. Thanki, Nicola Soranzo, Javier Herrero, Wilfried Haerty, Robert P. Davey

## Abstract

**Background:** Phylogenetic information inferred from the study of homologous genes helps us to understand the evolution of genes and gene families, including the identification of ancestral gene duplication events as well as regions under positive or purifying selection within lineages. Gene family and orthogroup characterisation enables the identification of syntenic blocks, which can then be visualised with various tools. Unfortunately, currently available tools display only an overview of syntenic regions as a whole, limited to the gene level, and none provide further details about structural changes within genes, such as the conservation of ancestral exon boundaries amongst multiple genomes.

**Findings:** We present Aequatus, a standalone web-based tool that provides an in-depth view of gene structure across gene families, with various options to render and filter visualisations. It relies on pre-calculated alignment and gene feature information typically held in, but not limited to, the Ensembl Compara and Core databases. We also offer Aequatus.js, a reusable JavaScript module that fulfils the visualisation aspects of Aequatus, available within the Galaxy web platform as a visualisation plugin, which can be used to visualise gene trees generated by the GeneSeqToFamily workflow.

**Availability:** Aequatus is an open-source tool freely available to download under the MIT license at https://github.com/TGAC/Aequatus. A demo server is available at http://aequatus.earlham.ac.uk/. A publicly available instance of the GeneSeqToFamily workflow to generate gene tree information and visualise it using Aequatus is available on the Galaxy EU server at https://usegalaxy.eu.

## Introduction

Sequence conservation across populations or species can be investigated at multiple levels from single nucleotides, to discrete sequences (e.g. transcription factor binding sites, exons, introns), genes, genomic blocks, and chromosomes. Analyses at each of these levels inform different evolutionary processes and time scales. While the vast majority of analyses focus on gene evolution, synteny, (the conservation of genomic blocks between multiple species) can be used to trace chromosome evolutionary history [1] and infer evolutionary relationships between genes across or within species [2]. Synteny resolution and analysis typically involves carrying out multiple sequence alignments (MSAs) and phylogenetic reconstruction, comprising multiple steps that can be computationally intensive even for relatively small numbers of data points [3].

Many methods are available for the identification of genome-wide orthology (MSOAR [4], OrthoMCL [5], OMA [6], HomoloGene [7], PhyOP [8], TreeFam [9], TreeBeST [10]). However, most of them do not incorporate taxonomic information (typically in the form of a species tree) while finding gene families, nor provide any information regarding transcript and protein structural changes across orthogroup members. The Ensembl GeneTrees pipeline [11], a computational workflow developed by the EMBL-EBI Ensembl Compara team, produces familial relationships based on clustering, MSA, and phylogenetic tree inference. The gene trees in Ensembl Compara are inferred with TreeBeST, which relies on a reference species tree to guide the process and calculates the probability of a gene tree in the context of species evolution. The data are stored in a relational database which contains information on gene families, syntenic regions and protein families. In parallel, the Ensembl Core databases store gene feature information and other genomic annotations at the species level. The Ensembl project (release 90, August 2017) at EMBL-EBI houses 100 vertebrate species [12], along with precomputed MSAs and gene family information.

Phylogenetic reconstruction is the most traditional method to represent and view comparative datasets across a given evolutionary distance, but specific tools such as Ensembl Browser [13], Genomicus [14], SyMAP [15], and MizBee [16] also exist to provide finer-grained information. These tools are able to provide an overview of syntenic regions as a whole, with only Genomicus reaching down to the gene order and orientation level. Conversely, phylogenetic trees retain ancestral information but do not represent the underlying information regarding structural changes within genes, such as the conservation of ancestral exon boundaries between multiple genomes or variants within genes that can be correlated to phenotypic changes. In order to build these gene-level visualisations, basic genomic feature information is required.

Therefore, we have developed Aequatus to bridge the gap between phylogenetic information and gene feature information. Here we show that Aequatus allows the identification of exon/intron boundary changes and mutations, informing the user about underlying genetic changes, but can also highlight mis-annotations, pseudogenes [17], or polyploidisation in animal and plant genomes.

## Materials and Methods

Aequatus is built using open-source technologies and is divided into a typical server-client architecture: a web interface and a server backend (see Figure 1).

The server-side component is implemented using the Java programming language. It retrieves and processes comparative genomics information directly from Ensembl Compara and Ensembl Core databases. Pre-calculated gene trees and genomic alignments, in the form of CIGAR strings [18], are held in Ensembl Compara, which are cross-referenced by Aequatus to Ensembl Core databases for each species to gather genomic feature information using the unique gene stable IDs.

**Figure 1:**
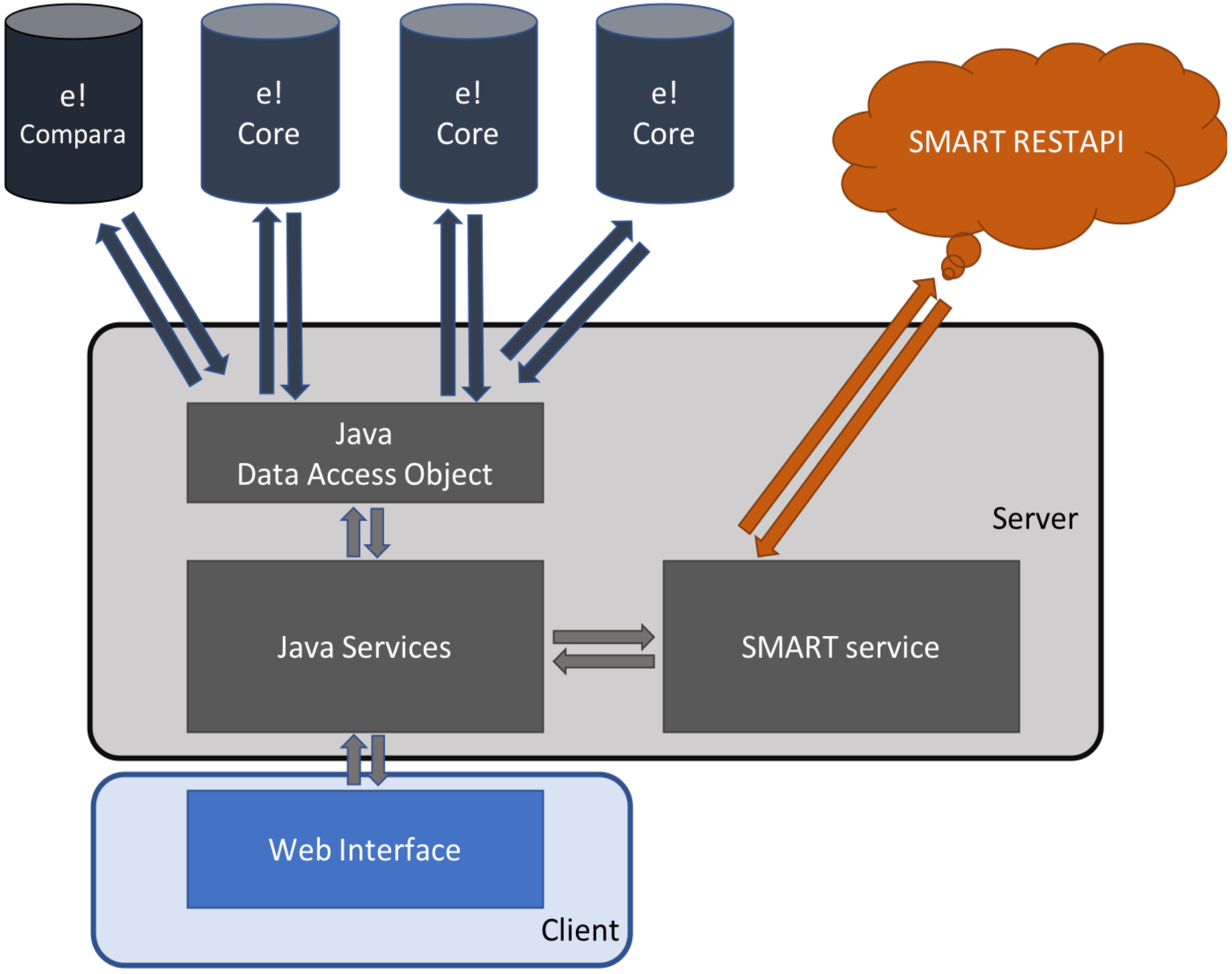
The Aequatus infrastructure, showing the interactions between the server-side implementation, connected to Ensembl compara and core database using Java Data Access Objects and SMART server via REST API, and the client-side implemented using popular techniques such as JavaScript, jQuery, d3.js and jQuery DataTables.

The Aequatus web interface comprises well-known web technologies such as SVG, jQuery, JavaScript and D3.js [19] to provide a fast and intuitive web-based browsing experience over complex data. Comparative and feature data are processed and rendered in a intuitive graphical interface to provide a visual representation of the phylogenetic and structural relationships among the set of chosen species.

Aequatus visualises gene families using a phylogenetic tree generated from gene sequence conservation information, held in a Ensembl Compara database, and gene features from Ensembl Core database. Gene features are presented in the form of exon-intron boundaries and 5’ and 3’ UTR. In this gene tree view, users are able to select a gene from a given species as a “guide gene”, and the homologous genes discovered through the comparative analysis are shown with respect to this guide gene. The representation of internal similarity among homologues is achieved by comparing the CIGAR strings for homologous genes with the CIGAR of the guide gene and mapping back to the homologous gene structure.

Aequatus is also able to visualise homologous genes in a customised Sankey view, using the d3.js [19] visualisation library, and provides feature information in an interactive Tabular view, using the jQuery DataTable [20] library. Statistical information for each member in a set of homologues, such as percentage coverage, positivity and identity, are fetched from homology and homology_member tables of the Ensembl Compara database.

We have integrated a SMART (Simple Modular Architecture Research Tool) [21] service to search for and visualise domain information of a protein sequence. We use the SMART REpresentational State Transfer (REST) API to retrieve protein domains, motifs, signal, repeats information from the SMART server using protein sequences.

Finally, to complement these various visualisations for the homologous genes and their gene trees, Aequatus provides gene order information in the form of a syntenic view (see Section 3). For a selected gene, homologues are fetched from homology and homology_member tables of the Ensembl Compara database. The neighbouring genes for these homologous genes are retrieved from the Ensembl Core databases using positional information and organised into a syntenic representation. Much like the shared conserved exon depiction in the gene tree view, syntenic genes are coloured based on the shared homology.

## Results

The landing page of Aequatus (see Figure 2) contains a header with a search box (2A) and a dropdown list of species (2B), followed by a selectable Chromosomal view underneath (2C).

**Figure 2:**
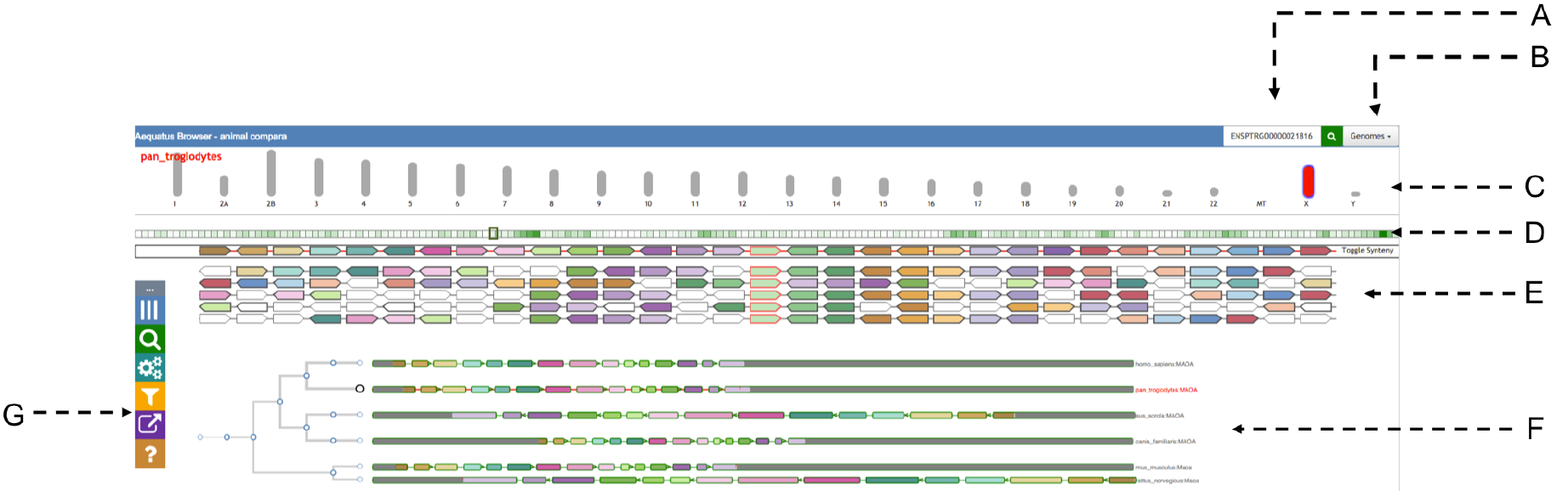
The main view of Aequatus. The header on top provides a search box (A) and a genome list (B). It is followed by the Chromosomal view (C), where the selected chromosome is coloured in red. Below there is an overview of genes (D) for the selected chromosome, followed by a zoomed area of the chromosome with genes shown in the syntenic view (E), and by the gene tree view (F). The Aequatus control (G) panel is visible on the far left.

Aequatus has a draggable control panel (2G) on the left-hand side, which contains buttons to show/hide the chromosome selector on top, modify gene views and labels, access to the search box and the export options, as well as a link to the help pages.

### 1. Aequatus user interface

Aequatus provides various ways to visualise gene trees and the inferred orthology/paralogy from them.

### 1.1 Main Gene Trees View

The gene tree view (see Figure 3) comprises a phylogenetic tree on the left, built from GeneTree information stored in a Ensembl Compara database [11]. Aequatus relates the genes through different events (e. g. duplication, speciation, and gene split) for the gene family and homologous genes against each respective node, which are coloured based on the potential evolutionary event. The selected guide gene is depicted as a larger circle black leaf node in the tree, with a red label on the right, while the other genes have a smaller circle leaf node and a grey label.

**Figure 3.**
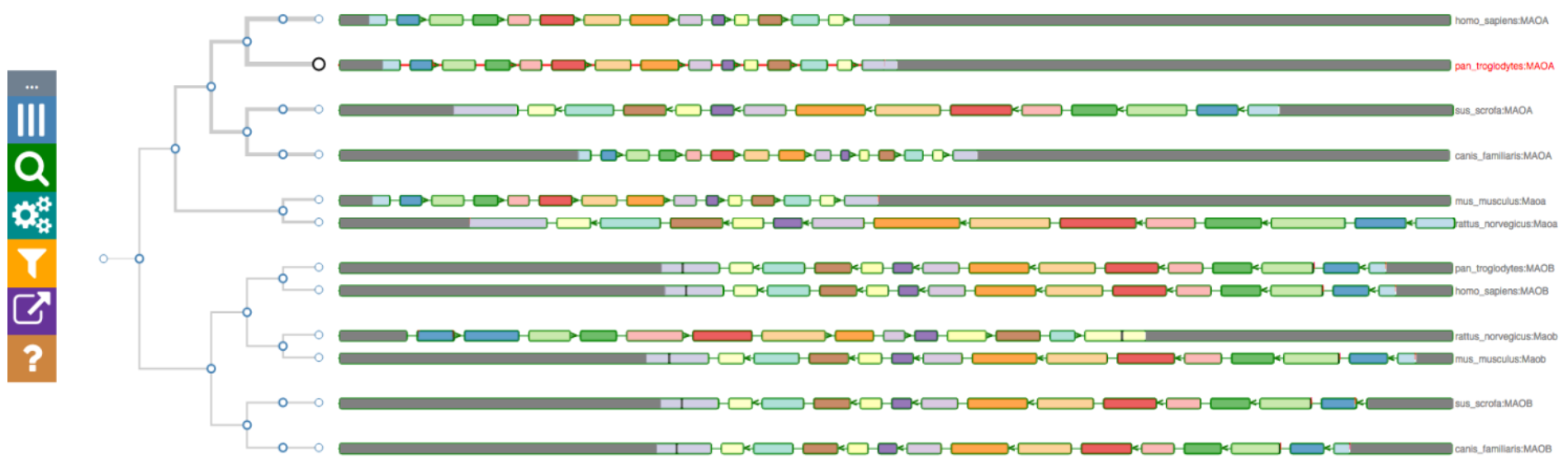
The gene tree for the monoamine oxidase (MAO) gene, with the pan_troglodytes gene as the reference, alongside other homologous genes in the exon-focused view. Considering the gene tree on the left, it is clear that the MAO genes are separated into two clusters, corresponding to the MAO-A and MAO-B gene families.

On the right, Aequatus depicts the internal gene structure, using a shared colour scheme for coding regions, to represent similarity across homologues. Homologous genes are visualised by aligning them against a given guide gene. Aequatus is also able to indicate insertions and deletions in homologous genes with respect to shared ancestors. Black bars within exons represent insertions, while red lines represent deletions specific to a given gene compared with the reference.

Aequatus provides two view types for gene families. The first (default) view is exon-focused (as in Figure 3), where all introns are set to a fixed width, since long introns can adversely affect the visibility of surrounding exons. This provides easier browsing of the actual gene structure, especially when less screen real estate is available. Conversely, in the second view all homologous genes are resized to the maximum available width in the web browser, showing introns and exons proportional to the real gene size. Users can switch between these views from the “Introns” settings in the control panel.

### 1.1.1 Popups

Aequatus provides a contextual menu system via interactive popup menus, which are displayed when a user clicks on a gene (see Figure 4). Each popup shows: the gene name and its position; a link to find protein domain information using SMART; links to export the protein sequence or the CIGAR alignment; an option to set the current gene as the guide in order to see insertions and deletions in homologous genes relative to the selected guide gene; a link out to the Ensembl page for the gene; an option to view the pairwise alignment.

**Figure 4.**
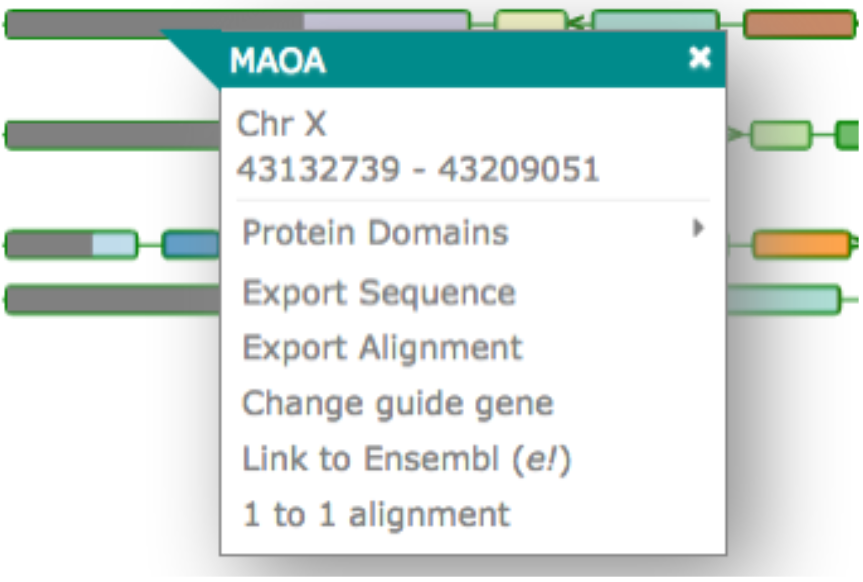
The popup in the gene tree view when clicking on a gene. The popup contains the chromosome name and position, and options to view the protein domains, export the sequence or the alignment, change the guide gene, connect to the Ensembl page for the gene, and view the pairwise alignment.

### 1.1.2 Protein Domain

Aequatus can provide an interactive visualisation of the protein domains for the selected gene. Aequatus finds the protein domains by connecting to the SMART web server via its REST API and querying the protein sequence for domains, motifs, internal repeats, etc. In this view (see Figure 5), a user can filter and sort domains based on type, E-value, position and source of domain. The features shown in the diagram can be exported in CSV or Excel file format.

**Figure 5:**
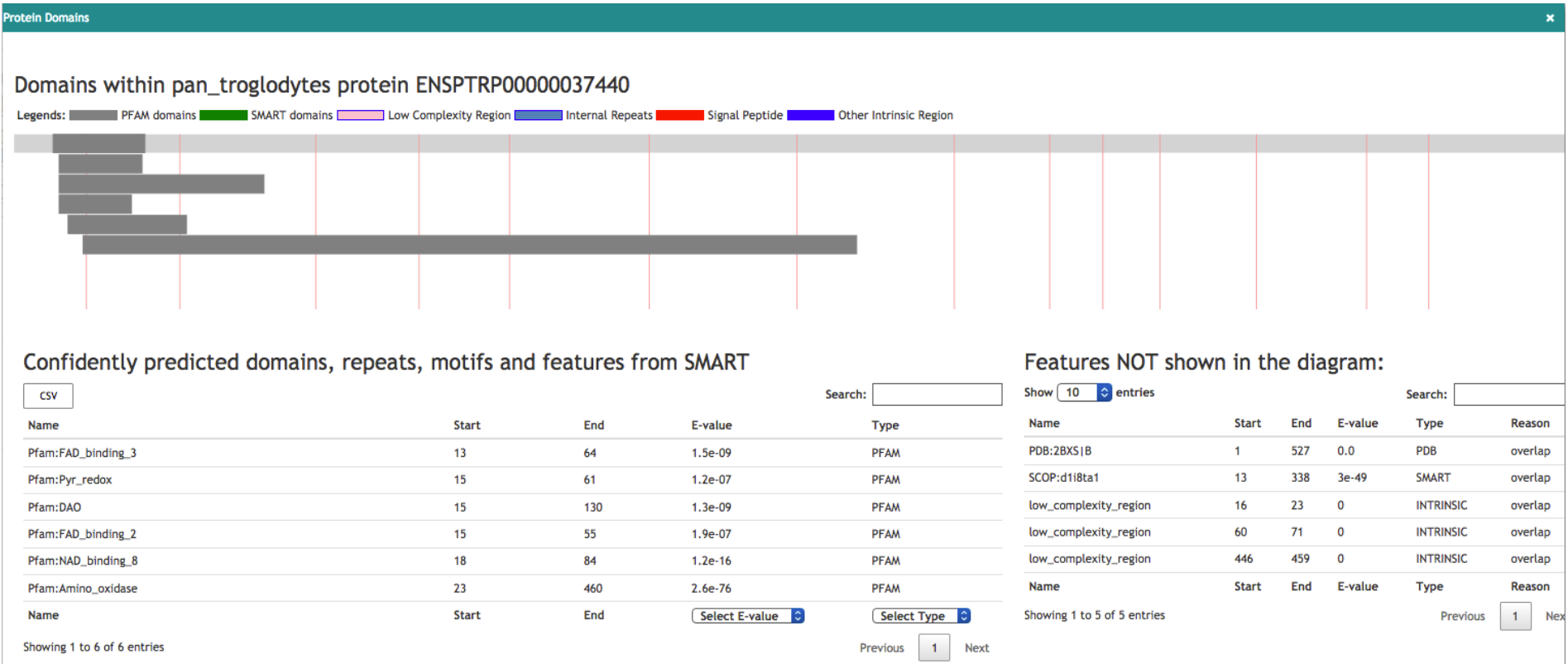
Visualisation of the protein domain information for the protein ENSPTRP00000037440 retrieved from the SMART server. On the top, drawings of domains mapped on exons. The tables below are listing the features shown in the diagram, as well as hidden features.

### 1.2 Homologous Genes

The underlying information describing homologous genes contained within the Compara database schema can be visualised using either a tabular view or Sankey plot.

### 1.2.1 Tabular View

The Tabular view (see Figure 6) contains statistical information for the homologous relationships. This view is dynamic, allowing the user to search for any homolog using a search box (6A) as well as filter results for the type of homology (6E) (1-to-1 orthologs, 1-to-many orthologs, and paralogs) or one or more specified species (6D). Homologous genes can be exported from the tabular view as Excel, CSV or PDF.

**Figure 6:**
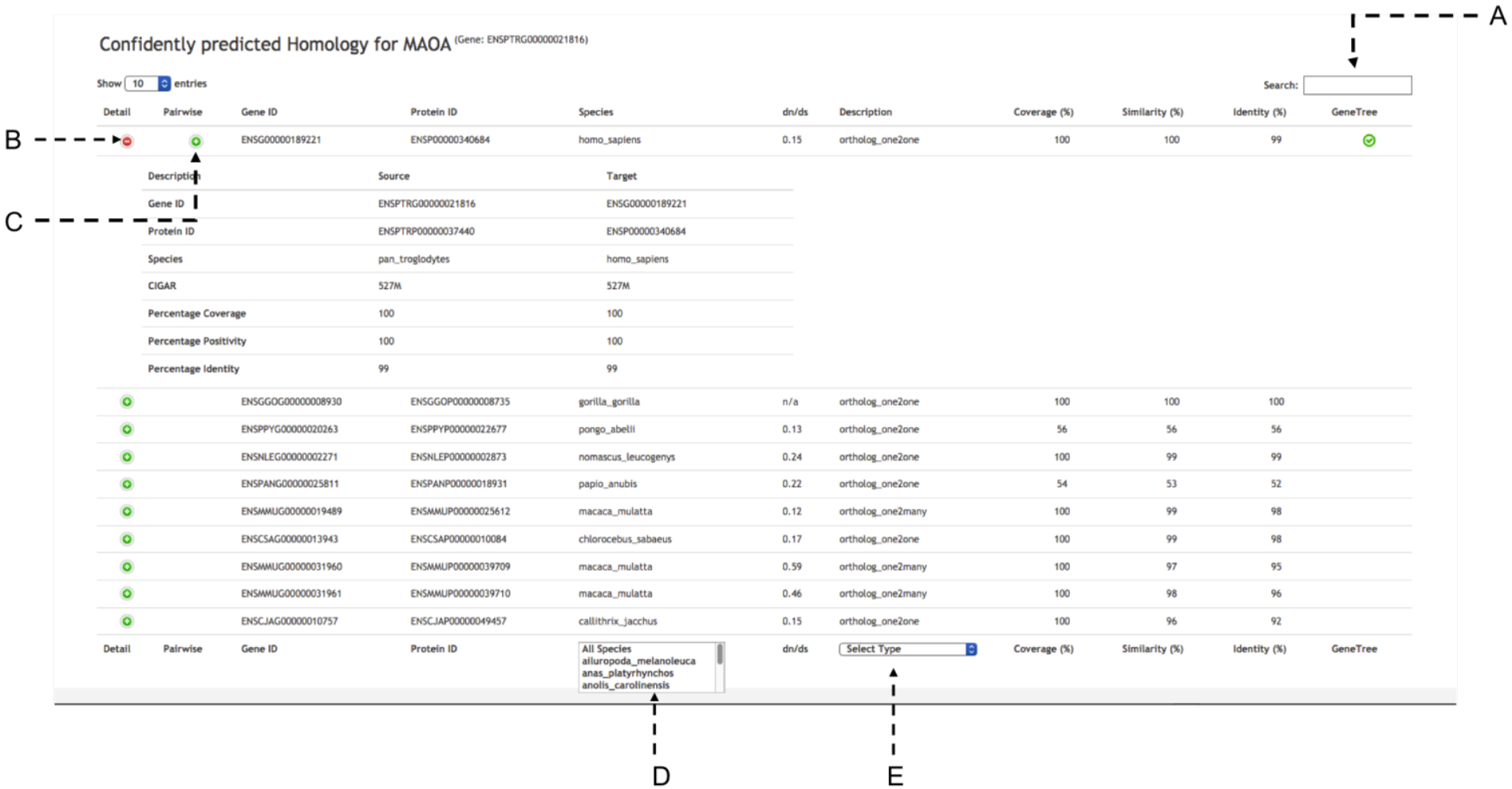
Homologous for the gene MAOA (ENSPTRG00000021816) in tabular view with statistical comparison about homologues. The tabular view contains a search box on top (A). There are 2 buttons to visualise statistical comparisons (B) and pairwise alignment (C) for each homolog. At the bottom it is possible to select from a list of species list (D) and the type of homology (E).

Extra details for the pairwise alignment between homologues can be shown by using the ‘+’ button for the homologue entry. The first button (6B) will show statistical comparisons for identity, coverage, similarity etc., while the second button (6C) will visualise the pairwise alignment with the gene structures as detailed below (Figure 8A).

### 1.2.2 Sankey view

The Sankey view (see Figure 7) visualises homology as a interactive diagram, where the homologues of a selected gene are distinguished by homology type, i.e. paralogs, 1-to-1 orthologs, or 1-to-many orthologs. The nodes for homologous genes are coloured by species, which helps finding genes from the same species in the case of 1-to-many and many-to-many orthologs.

**Figure 7:**
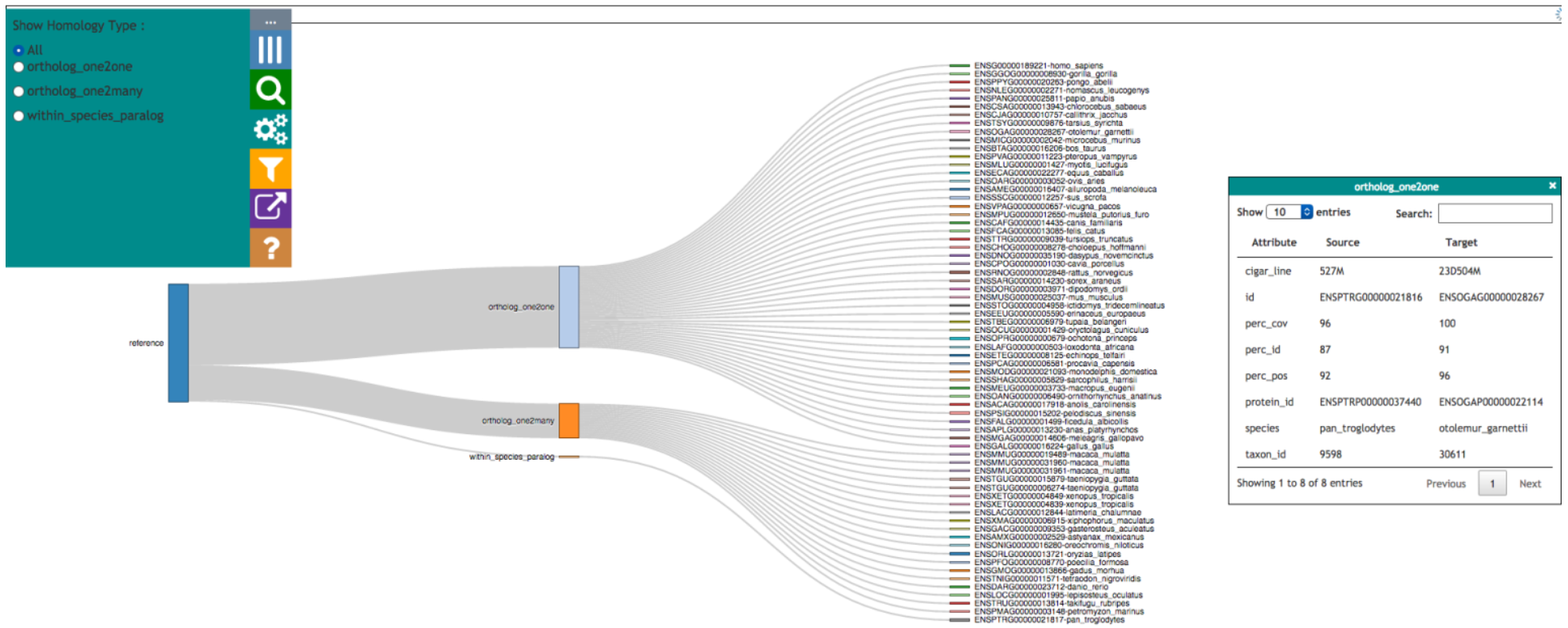
Homologues for a gene in Sankey format, grouped together by type of homology. The control panel on the left shows filters for the view. Further information for any homologue can be retrieved by clicking on it and is then shown in right box.

When clicking on a homologous gene, additional details for the homologous pair are displayed in the info panel on the right-hand side.

### 2. 1-to-1 alignment

1-to-1 alignments between homologous genes are important for pairwise comparison. 1-to-1 alignments (Figure 8) can be seen by clicking on the corresponding option either in the popup for the gene tree view or in the homologous genes tabular view. This will fetch the relevant alignment from the homology table of the Ensembl Compara database and visualise it based on the gene structure (8A), together with the pairwise protein sequence alignments (8B).

**Figure 8:**
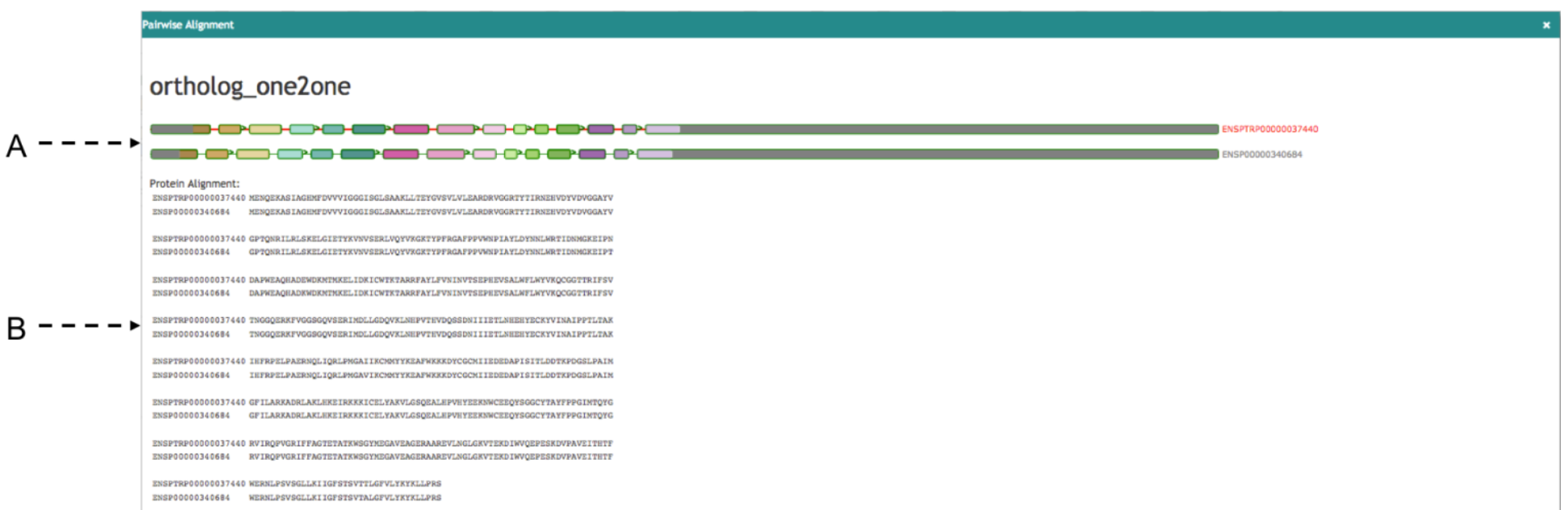
1-to-1 alignments between homologous genes. On the top (A) visualising alignment on gene structure and on the bottom (B) visualising pairwise sequence alignments.

### 3. Gene Order

Genes that share a common ancestor and are part of a consecutive block of genes are likely to have a transcriptional and/or functional relationship [22]. Hence, inferred homologues which are present in all species and in the same order are more likely to be real homologues. In the Gene Order view, neighbouring genes are displayed for the selected gene and its homologues (see Figure 9). Homologues of the genes in neighbouring species are coloured based on the matching genes from the reference species. Clicking on a gene feature will open a search panel with various viewing options, and mousing over a given gene will highlight all homologous genes within the same region. The syntenic view complements the main functionality of Aequatus by providing evidence for the conservation information for the genes of interest.

**Figure 9:**
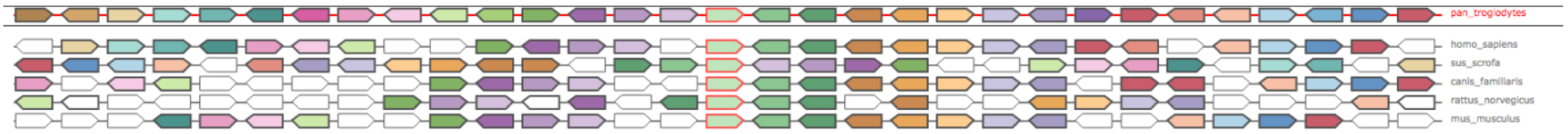
Gene Order for the MAOA gene in *Pan troglodytes*, where they are coloured by homologous genes. The selected gene and its homologous have a red border. White genes are the ones which don’t have any homologous in current visible region.

### 4. Search

Aequatus has keyword-based search functionality, whereby the user can provide search terms and a list of all the relevant genes is returned. Aequatus can query for matching gene symbols, Ensembl stable IDs (unique identifiers in the Ensembl project for each genomic annotation), common names for genes and proteins, or any keyword in the description. Search results then allow the user to visualise the corresponding gene tree, or homologous genes in the tabular or Sankey views.

### 5. Export

Users can export data at different points in the visualisation. In the gene tree view the underlying genomic data for the gene families can be exported in various forms, such as a list of gene IDs, the sequence alignments, or the gene trees in Newick [23] or JavaScript Object Notation (JSON) [24] format, for use in downstream tools. The tabular view can be exported in CSV, XLS, and PDF format.

### 6. Persistent URLs

Aequatus provides persistent unique URLs to enable consistent access to genes of interest, making it easy to go back to the results of a previous search, to share information with collaborators, or for use in publications. Users can share the link for the visualisation of a specific gene, the results of a search for a term, or a specific reference to a given species and chromosome.

### Discussion

The ultimate goal of Aequatus is to provide a unique and informative way to render and explore complex relationships between genes from various species at a level of detail that has so far been unrealised in a single platform. While applicable to species with high-quality gold-standard reference genomes present in core database resources such as human or mouse, Aequatus has been designed to accommodate users that need to explore large, fragmented, non-model genome references that are held in institutional databases. We are testing Aequatus with a range of non-model organisms, such as koala, polyploid crops, and spiny mouse. As Aequatus can visualise relationships using simple CIGAR strings, any tool that outputs this format can use Aequatus to view them. We produce input for Aequatus using the GeneSeqToFamily pipeline, a freely available Galaxy workflow [25] for finding and visualising gene families for genomes which are not available from Ensembl databases.

In order to make Aequatus more accessible and reusable, the gene tree visualisation module from the standalone Aequatus browser is available as Aequatus.js [26], an open source JavaScript library. In this way, it preserves the interactive functionality of the Aequatus browser tool but can be integrated with other third-party web applications. We have demonstrated this by integrating the Aequatus.js library into Galaxy [27] as a part of the GeneSeqToFamily workflow.

### Future Directions

The main extension to the functionalities of Aequatus is the incorporation of Ensembl REST API functionality [28], where Aequatus will be able to retrieve information directly from Ensembl Compara and Core databases held at the EMBL-EBI, without any need for local database configuration. Whilst this will mean that users will need a reliable internet connection, it will reduce the need for local storage space for the Ensembl databases, improving the portability of Aequatus.

We also intend to containerise Aequatus using Docker and CyVerse UK [29], and BioConda [30] with Galaxy [25,27]. We will produce new APIs between Aequatus and TGAC Browser [31] to provide a comprehensive solution for genome analysis and exploration focused on non-model organisms.

## Acknowledgements

This work was strategically funded by the BBSRC (BBS/E/T/000PR5885, BBS/E/T/000PR9817) and through the EU TransPlant grant (BBS/E/T/000GP006). GeneSeqToFamily and the EI Galaxy platform are funded through the BBSRC-supported EI National Capability in e-Infrastructure (BBS/E/T/000PR9814).

This work was supported in part by the NBI Computing Infrastructure for Science Group, which provides technical support and maintenance to EI’s high-performance computing cluster and storage systems, which enabled us to develop this tool.

Conflict of Interest: none declared.

